# A novel approach for the extraction of nucleic acids using a hybrid paper-plastic device

**DOI:** 10.1101/2025.09.29.678505

**Authors:** Sowjanya Goli, Umarani Brahma, Sachin Chandankar, Suman Karadagatla, Anamika Sharma, Vasundhra Bandari, Ira Bhatnagar, Amit Asthana

## Abstract

Nucleic acid–based diagnostics play a crucial role in early and accurate disease detection. However, conventional extraction approaches are expensive, time-consuming, and require complex instrumentation and skilled personnel, which restricts their use in resource-limited settings. To develop and evaluate a simple, low-cost, and equipment-free paper-based microfluidic cassette device for rapid extraction of nucleic acids (DNA and RNA) from biological samples. A paper–plastic cassette was fabricated by laminating filter paper or glass fiber substrates between thermally bonded plastic sheets. The system was tested using Klebsiella pneumoniae (DNA) and HeLa cells (RNA) with different lysis buffer formulations. Extracted nucleic acids were quantified using a Qubit fluorimeter and assessed by gel electrophoresis and PCR amplification. Results were compared with standard commercial extraction kits. Among tested substrates, LF1 combined with lysis buffers 2, 4, and 5 yielded the highest quality DNA and RNA with 260/280 absorbance ratios of approximately 1.8 (DNA) and 2.0 (RNA). The nucleic acid yields were comparable to commercial kits, and the extracted material was suitable for direct downstream applications, including PCR. The developed paper–plastic cassette enables rapid, affordable, and equipment-free extraction of nucleic acids. This platform shows strong potential for point-of-care molecular diagnostics, particularly in low-resource and decentralized healthcare settings.

## Introduction

Nucleic acid extraction is a foundational process in molecular biology and diagnostics [1]. It is essential for applications such as polymerase chain reaction (PCR), loop-mediated isothermal amplification (LAMP), next-generation sequencing (NGS), and biosensor development [2]. Traditional methods for nucleic acid extraction, including phenol-chloroform precipitation, spin-column extraction, and magnetic bead-based separation, are time-consuming, require skilled personnel, and involve expensive instrumentation, such as centrifuges and pipetting robots [3]. These protocols are ill-suited for resource-limited settings and point-of-care (POC) applications, where access to laboratory infrastructure is limited [4]. Microfluidic technologies have emerged as promising solutions for miniaturizing and automating sample preparation, including nucleic acid extraction [5]. These systems allow fluid handling at the microliter or nanoliter scale, reducing reagent consumption and enabling integration with downstream analyses [6]. However, many microfluidic devices are fabricated using polydimethylsiloxane (PDMS) or thermoplastics and rely on external tubing, syringe pumps, or pressure controllers for fluid actuation [7]. The dependence on such peripherals compromises the portability and field deployability of these systems, particularly for use in decentralized environments [8]. Paper-based microfluidic devices (μPADs) have gained prominence for overcoming the limitations of conventional and PDMS-based microfluidics owing to their simplicity, low cost, biocompatibility, biodegradability, and pump-free operation using capillary flow [9]. Since the introduction of μPADs by Whitesides et al. in 2007, μPADs have been successfully used in a range of diagnostic applications, including glucose testing, heavy metal detection, and immunoassays [10]. The porous nature of cellulose-based substrates allows for spontaneous fluid movement without external power sources, making them ideal candidates for sample-to-answer systems in low-resource settings [11]. Despite their potential, μPADs have not yet been fully optimized for complex molecular workflows, such as nucleic acid extraction [12]. Existing approaches often incorporate tubing for sample injection or external heaters for lysis, which compromises the inherent portability and simplicity of paper-based systems [8]. Moreover, some μPADs embed chemical reagents directly into the device, but often lack sequential control over reagent delivery or efficient binding matrices for nucleic acid capture and purification [13]. Recent efforts have focused on integrating nucleic acid extraction steps into paper devices using silica-modified regions, chaotropic salts for cell lysis, and wax-patterned flow channels to control fluid sequencing. However, many of these methods still require multistep manual operations or are limited to proof-of-concept demonstrations. In Addition, the inclusion of external tubes or elution reservoirs reintroduces the complexities that μPADs aim to eliminate [14].

In this context, we propose a novel, fully tubeless, paper-based microfluidic device designed for efficient nucleic acid extraction from biological fluids such as saliva, urine, and blood. The system relies solely on capillary action to drive fluid flow across various predefined hydrophilic and hydrophobic zones, enabling sequential execution of lysis, washing, and elution steps without the need for pumps, valves, or tubing. This innovation is based on its structural simplicity and functional versatility: 1) It integrates dry stored reagents for on-demand activation during the fluid contact. 2) It uses a silica-modified capture zone embedded in cellulose for selective binding of nucleic acids. 3) This ensures flow directionality and timing through a tailored channel geometry and wax barriers. Our device supports field-ready, low-cost, and power-free nucleic acid isolation in under 15 minutes, and the eluted nucleic acids are directly compatible with isothermal amplification methods, such as LAMP, and colorimetric detection, without requiring cold-chain logistics or further purification.

## Materials and Methods

Whatman qualitative filter paper grade 4 with a thickness of 210 µm and retaining particles measuring 25 µm, LF1 glass fiber with thickness of 247 μm, wicking rate of 35.6 s/4cm, and water absorption rate of 25.3 mg/cm^2^, and VF2 glass fiber with thickness of 785 μm, wicking rate of 23.8 s/4 cm, and water absorption rate of 86.2 mg/cm^2^ were purchased from Cytiva Life Sciences (USA). Ammonium Chloride, Potassium bicarbonate, EDTA, were purchased from (SRL Pvt. Ltd Hyderabad) Diethylpyrocarbonate (DEPC), Bought from (Sigma Aldrich, Hyderabad) Tris Base, Hydrochloric acid (HCL), Boric Acid, Guanidine Hydrochloride, Tris Hydrochloric Acid, Triton X-100, Guanidine Isothiocyanate, Sodium Dodecyl Sulphate (SDS), Ethanol, were procured from (Himedia Pvt. ltd, Mumbai) Lithium Chloride, Tween 20, Tris buffer, Sodium Chloride, Ethidium Bromide, TAE, Autoclaved MiliQ, Agarose, DNA loading dye, DNA ladder, (Promega) Taq polymerase, 1OmM mg+2 buffer, dNTP, 16s Ribosomal primers.

### Design and Fabrication of Paper-Plastic Device

The device was designed using Laser CAD software and was fabricated using a paper substrate. The paper substrate was cut into circular discs with a diameter of 8 mm. These discs were positioned between two thermally bonded lamination sheets, each pre-cut into a 15 mm × 15 mm square. A circular window with a diameter of 7 mm was cut into both lamination sheets to expose the paper substrate from both sides, enabling dual-sided fluid interaction. The design was cut using an SIL computer-controlled laser-engraving system. For the final assembly, the paper disc was aligned within the cut-out regions of the lamination sheets and thermally sealed using a laminator (Vision TEK Laminator). The assembly results in a compact, sealed, paper-plastic device.

### Sample Preparation and Nucleic Acid Extraction

Standard bacterial isolates of Klebsiella pneumoniae were used to evaluate the performance of the system using biological samples. The bacterial cultures were incubated overnight in Tryptone Soya Broth (TSB; HiMedia, Mumbai) at 37□°C with shaking at 150 rpm. Before genomic DNA isolation, the cells were centrifuged at 4000xg for 10 min and washed with 1x Phosphate buffer saline. Genomic DNA was isolated using both the developed system and the Promega nucleic acid extraction kit (used as a control) following the manufacturer’s guidelines for comparative analysis. HeLa cancer cells were used to investigate further the adaptability of the system across different biological sources. The cells were cultured in high-glucose Dulbecco’s Modified Eagle’s medium (DMEM; HiMedia, Mumbai, India) supplemented with 10% fetal bovine serum (FBS). The cultures were maintained in plastic culture dishes at 37□°C in a humidified atmosphere containing 5% CO□. Following the culture, RNA was extracted using both the developed system and TRIzol extraction method, following the manufacturer’s guidelines for comparative analyses. After RNA isolation, samples were treated with proteinase K and DNase I to remove protein contaminants and genomic DNA. Both DNA and RNA were quantified using a Qubit fluorimeter (Thermo Scientific, USA), and 500 ng per sample was run on a 0.8% agarose gel electrophoresis.

PCR amplification (Bio-Rad, Thermocycler) was performed to determine the yield and quality of the extracted nucleic acids. Control DNA and device-extracted DNA were used as PCR templates. DNA amplification was performed using 10 μL of the extracted DNA in a 25 μL reaction mixture containing 10× PCR buffer (NEB), 10 mM of each dNTP, 0.5 μL of each primer, and 0.12 μL Taq DNA polymerase (NEB). The 16S rRNA amplification was conducted with an initial denaturation step at 94□°C for 5 min. The amplification cycle consisted of 30 cycles of 95□°C for 15 s, annealing at 59□°C for 1 min, extension at 72□°C for 1 min, and a final elongation step at 72□°C for 10 min.

After RNA extraction, RNA was converted to cDNA using the PrimeScript 1st Strand cDNA Synthesis Kit (Takara), according to the manufacturer’s instructions. The cDNA was used as a template to confirm the amplification of β-actin. Amplification was conducted using an initial denaturation step at 95°C, followed by 35 cycles of annealing at approximately 60°C, extension at 72°C, and a final extension step at 72°C. The amplified products were analyzed by agarose gel electrophoresis using a 2% gel prepared with ethidium bromide (EtBr) as an intercalating dye. The gels were imaged and documented using a GelDoc visualization system (Bio-Rad).

### The composition of the lysis and wash buffers for them accordingly

Lysis buffer 1 (L□) consisted of 5.5 M Guanidine hydrochloride, 50 mM Tris-HCl, 20 mM EDTA, and 1.3% Triton X-100. Lysis buffer 2 (L□) contained 5.5 M Guanidinium isothiocyanate, 100 mM Tris-HCl, 10 mM EDTA, 10% Triton X-100, and 1% sodium dodecyl sulfate (SDS). Lysis buffer 3 (L□) was composed of 100 mM Tris-HCl, 10 mM EDTA, 10% Triton X-100, and 1% sodium dodecyl sulfate (SDS). The wash buffers used for L1, L2, and L3 were identical. Wash buffer 1 consisted of 70% ethanol, whereas wash buffer 2 consisted of 100% ethanol. Lysis buffer 4 (L□) contained 7.2 M LiCl, 5% Tween 20, 5% Triton X-100, 10 mM EDTA, and 50 mM Tris-buffered saline. The wash buffers for L4 included wash buffer 1, which was composed of 5 M LiCl and 55% ethanol. Wash buffer 2 consisted of 5 mM EDTA, 70% ethanol, and 100 mM Tris-HCl, with the pH adjusted to 7.6. Lysis buffer 5 (L□) contained 6 M Guanidine hydrochloride, 1% Tween 20, and 1% sodium dodecyl sulfate (SDS). Lysis buffer 6 (L□) was composed of 4.5 M Guanidine hydrochloride, 50 mM Tris-HCl, and 30% Triton X-100. The wash buffers used for L4 and L5 were identical. Wash buffer 1 consisted of 5 M Guanidine hydrochloride, 20 mM Tris HCl, and 40% ethanol, with the pH adjusted to 6.6. Wash buffer 2 was composed of 50% ethanol, 20 mM NaCl, and 2 mM Tris-HCl, with the pH adjusted to 7.5.

### Device optimization and evaluation for extraction

The design and performance of the paper–plastic microfluidic cassette were systematically optimized to develop a portable, equipment-free platform for nucleic acid extraction. The paper disc was 8 mm in diameter, allowing it to retain up to 50 μL of the solution without leakage. The plastic lamination pouch featured a 7□mm central window, ensuring a tight peripheral seal when sandwiching the paper disc. These dimensions are critical for maintaining the device integrity and efficient fluid distribution during the extraction process. The laser cutting parameters were optimized to ensure precision without thermal damage. For paper substrate cutting, the maximum power was set to 20□W, and the minimum power was set to 10□W. For the lamination sheets, the maximum power was adjusted to 30W. In contrast, the minimum power was maintained at 10

W. An absorbent pad (150 × 300 mm) was placed beneath the device during operation to capture excess fluid and prevent contamination. To assess the nucleic acid-binding efficiency of the device, three paper substrates (Whatman Grade 4, LF1, and VF2) were coated with 10□μL lysis buffer (buffers 1–6), each targeting cell lysis and nucleic acid stabilization. After air-drying of the coated substrates, nucleic acid extraction was performed using an assembled cassette. The performance of each substrate buffer combination was evaluated based on the yield and purity of the extracted nucleic acids, which are essential parameters influencing the sensitivity and specificity of downstream molecular diagnostic techniques such as PCR. Among the tested configurations, optimal substrate–buffer pairs were identified that supported efficient extraction under ambient conditions, demonstrating the potential of the device as a tubeless, field-deployable nucleic acid purification system.

### Nucleic acid Extraction from the Sample

Different lysis buffers with related wash buffers were prepared, which could be used for solid-phase nucleic acid extraction. All buffers contained a detergent, or a combination of detergents, chaotropic reagents, nucleic acid complexing salts, and a chelator. Specific wash buffers were used to wash the device. Two wash buffers were used for each lysis buffer. The first wash buffers contained RNA complexing salt or chaotropic reagent at low concentrations compared to the respective lysis buffer, and the other wash buffers were composed of alcohol and chelator. All buffers were treated at the same volume.

### Protocol for the extraction of nucleic acids using a paper-plastic device

The laminated devices were saturated with lysis buffer and air-dried. Ten microliters of the sample were mixed with 20 μL of lysis buffer. The cocktail was then spotted on the pre-treatment. The discs were then washed with the respective wash buffers. Forty microliters of wash buffer and 80 μL of wash buffer 2 were used. An excess wash buffer was applied using an absorbent pad. After washing, the device was dried and eluted using 20 μL nuclease-free water or DEPC-treated water. The eluate was directly used as a template for cDNA synthesis and was later used for PCR amplification to confirm the presence of RNA. For DNA quantification, samples were directly subjected to agarose gel electrophoresis. Nucleic acids, DNA, and RNA were quantified using a Qubit fluorimeter.

### Sample volume optimization for extraction

Optimization of biological test samples is required. The gold standard method for nucleic acid extraction from the blood was used to optimize the volume needed for further reactions. The presence of DNA was confirmed by Qubit fluorimeter quantification, agarose gel electrophoresis, and Polymerase chain reaction, whereas the presence of RNA was confirmed by cDNA synthesis and PCR amplification. A series of nucleic acid extractions was performed with varying sample volumes, scaled down from 1000 μL to 10 μL. Based on these experiments, the minimum volume of the sample that could be processed on a paper device was determined to be 10μl.

## Results and Discussion

### Design of Paper-plastic cassette device

A paper-based microfluidic device was fabricated using two components: a paper-based substrate, where the specimen was loaded for processing, and a cassette casing made of a plastic sheet around the device for handling. An image and illustration of the device and its assembly are shown in Figure 1. A paper-plastic device was fabricated using a laser cutter and laminator. To facilitate the selection of the appropriate substrate material, this study compared the performance of three paper-based materials: LF1, VF2, and Whatman grade 4 filter paper. The physical and chemical properties of these paper-based materials were first investigated, and their NAE performance was observed. The extraction efficiencies of LF1, VF2, and Whatman grade 4 in combination with six types of lysis buffers were compared. To further understand the physical properties of the six paper-based NAE, the quantity, thickness, tightness, and hydrophilicity of these materials were investigated. Observations of Whatman grade 4 and VF 2 revealed that their thickness and high absorption capacity inhibited proper washing of the device. Compared to Whatman grade 4, LF1 had more uniform fibrils, which also facilitated the appropriate washing and wicking of liquids. The presence of glass membranes in LF1 enhanced nucleic acid binding.

**Figure 1:**
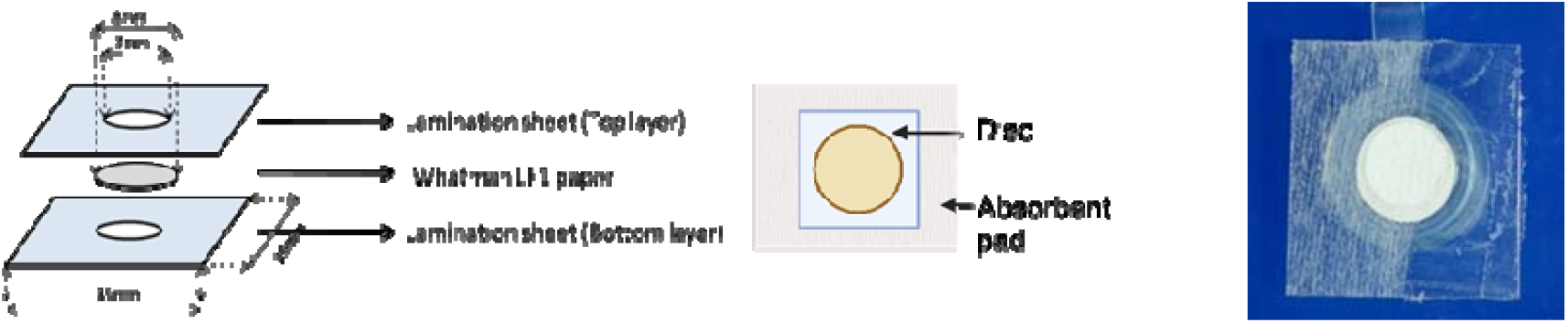
A. Illustration of paper-plastic device assembly; B. Image of paper-plastic device.

### Optimization of the device size

The disc size was optimized to 8 mm in diameter to contain 50 µL of solution without appreciable device leakage. For the plastic pouch, the opening was optimized to a 7 mm diameter to sandwich the paper disc properly without any gap on the periphery. Subsequently, the working parameters of the laser cutter device were optimized to prevent charring, with the maximum power set at 20 watts and the minimum power at 10 watts. Additionally, the working parameters of the laser cutter were optimized to a maximum power of 30 watts and a minimum power of 10 watts. The dimensions of the absorbent pad were 150 mm × 300 mm.

### Coating of the paper devices

To develop an easy nucleic acid purification technique that doesn’t depend on advanced lab equipment, we initially examined various chemicals for their potential to aid in capturing negatively charged DNA and RNA by spotting them onto paper substrates such as paper-plastic LF1, VF2, and Whatman grade 4. The substrates were coated with 10 μL of lysis buffers 1, 2, 3, 4, 5, and 6, with one lysis buffer for each. After the lysis buffers were successfully coated onto the paper device, the devices were air-dried before extraction. To identify the best paper substrate and suitable lysis buffer for nucleic acid extraction, three types of paper substrates were compared with six different lysis buffers and their respective wash buffers. Whatman filter paper Grade 4, LF1, and VF2 were used for nucleic acid extraction. The physical and chemical properties of the paper substrates were investigated, and their ability to extract nucleic acids was evaluated.

### Extraction of Nucleic Acids

10 μL of the biological sample was combined with 20 μL of lysis buffer and incubated for 10 minutes. Subsequently, the resultant lysate was introduced into the pre-treated device using the same lysis buffer.

### DNA extraction using a paper-plastic cassette

A 10 μL aliquot of the cultured Klebsiella pneumoniae sample with A random concentration was used as the standard. DNA was automatically extracted and purified with this device. To compare, genomic DNA was also extracted from 1 mL of bacteria at a specific concentration using the Promega Nucleic Acid Extraction Kit, a standard commercial method. The levels of concentration and purity across the three groups. LF1 (1-6), VF2 (1-6), and Grade 4 (1-6) were measured via absorbance with a Qubit fluorimeter. Results indicated that DNA concentration from the paper device was similar to that from the commercial method. DNA purity was assessed by the 260/280 absorbance ratio, with the average ratios for the samples obtained by different methods being 1.8.

The DNA extracts collected from our device had better purity than those obtained using other methods. This improvement was attributed to the specificity of the device and the limited number of steps required to prevent contamination from the external environment. The extracted DNA was used as a template for downstream PCR analysis to demonstrate the potential of this device for molecular applications. Although the efficiency was a bit lower than traditional methods, it still met the requirements for downstream applications. As shown in Figure 2, Qubit results demonstrated that the yield extracted from LF1> WF4>VF2. DNA retention on the paper substrate affects capture efficiency. The LF1 filter substrate was the best choice because of its hydrophilic properties, fast wicking speed, and high compatibility with PCR amplification. The significant differences in the flow rates of the three substrates also affected the efficiency of DNA extraction.

**Figure 2:**
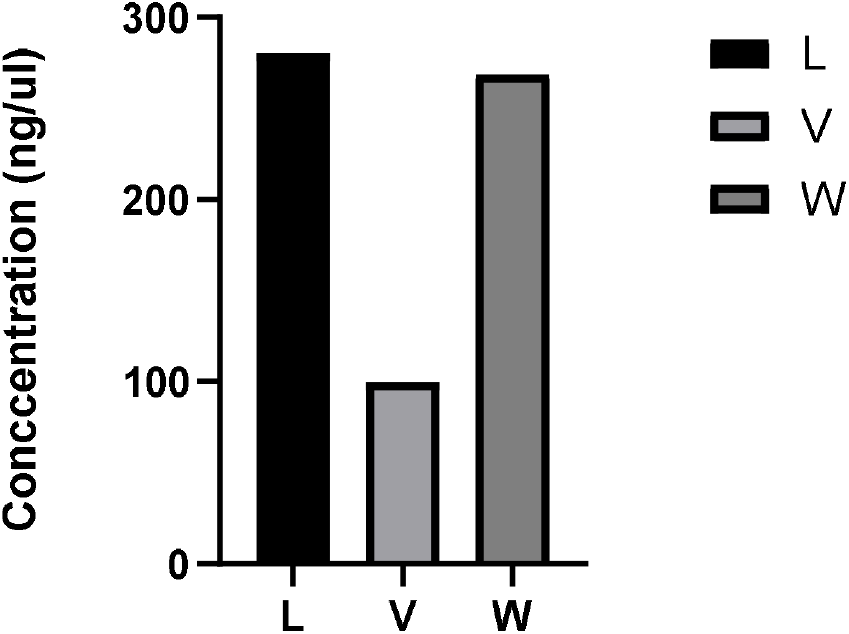
Comparison of DNA extraction yields between LF1, VF2, and W4. The figure shows the Qubit analysis results of the DNA samples.

**Figure 3:**
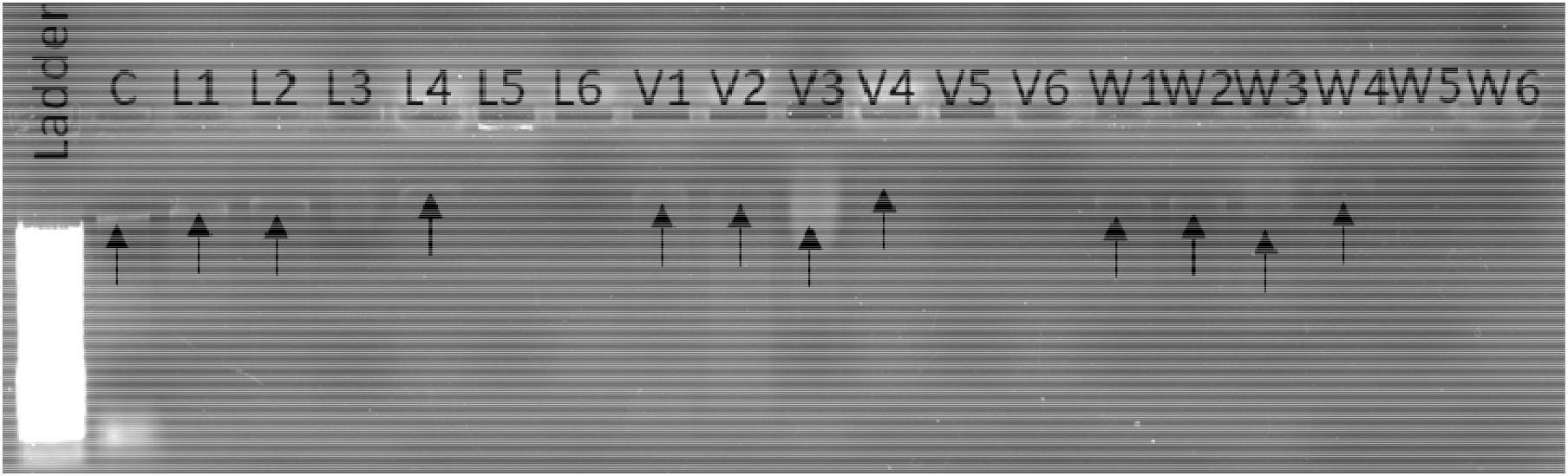
Electrophoretic analysis of DNA samples. DNA was extracted from a paper-plastic device. It is interpreted from the image that L1, L2, L4, L5, V1, V2, V3, V4, W1, W2, W3, W4 samples gave positive results for presence of DNA.

**Figure 4:**
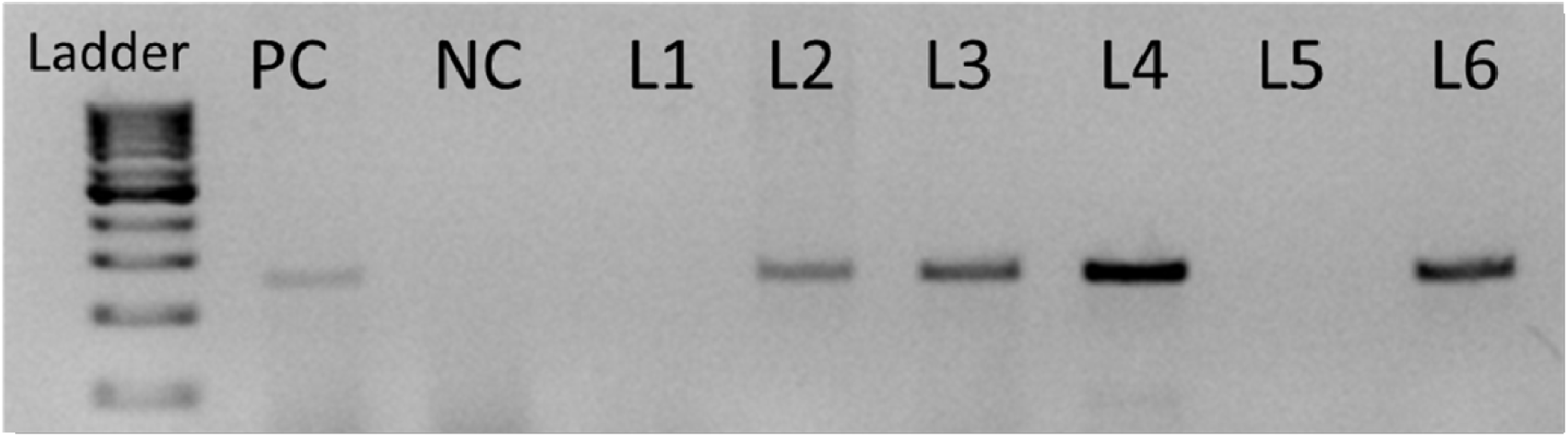
Electrophoretic analysis of PCR products with different paper devices. Checking the quality of DNA. It is interpreted from the image that L2, L3, L4, and L6 samples gave positive results for downstream applications like PCR.

### RNA extraction using a paper-plastic cassette

We further evaluated the versatility of the paper-plastic cassette for RNA extraction from HeLa cells. Ten microliters of cultured cells at A particular concentration were used as the standard, and RNA extraction was carried out using a paper device. To compare with traditional RNA extraction methods, RNA was extracted from 1 mL of cells at a consistent concentration using the TRIzol method. The concentration and purity of the RNA samples were quantified by measuring absorbance using a Qubit fluorimeter. The data showed that the concentration of RNA extracted from the paper device was similar to that of RNA obtained through the standard commercial method. The RNA purity was evaluated using the absorbance ratio at 260 and 280 nm. The average 260/280 ratio of the RNA samples obtained using different methods was 2 (Table. Of the three types of substrates, only the LF1 devices were able to extract pure RNA from the bacterial samples. To evaluate the RNA purity for downstream analysis, we conducted conventional PCR to amplify the actin housekeeping gene. Additionally, the RNA extracted from the system was amplified similarly to RNA obtained through the TRIzol extraction method. The LF1 device with buffers 1, 2, 4, and 5 combinations provided a positive yield for RNA.

**Figure 5:**
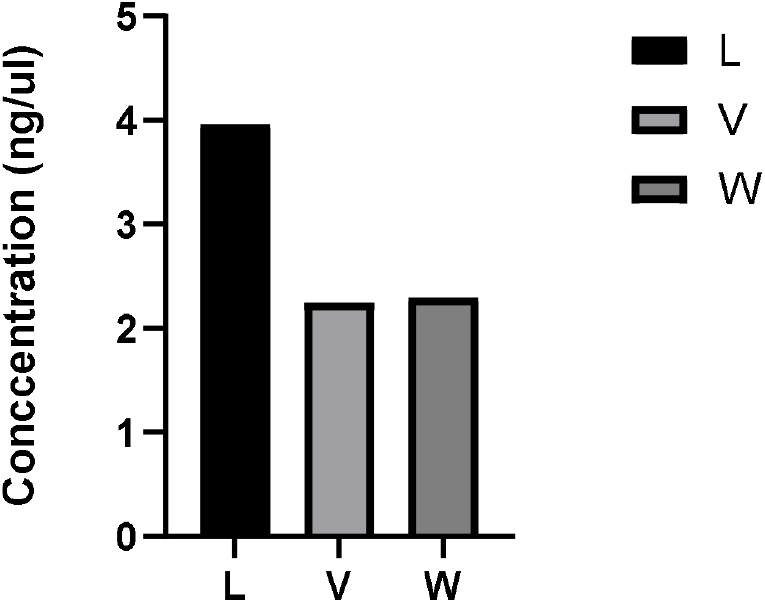
Comparison of RNA extraction yields between LF1, VF2, and W4. The figure shows the Qubit analysis results of the RNA samples.

**Figure 6:**
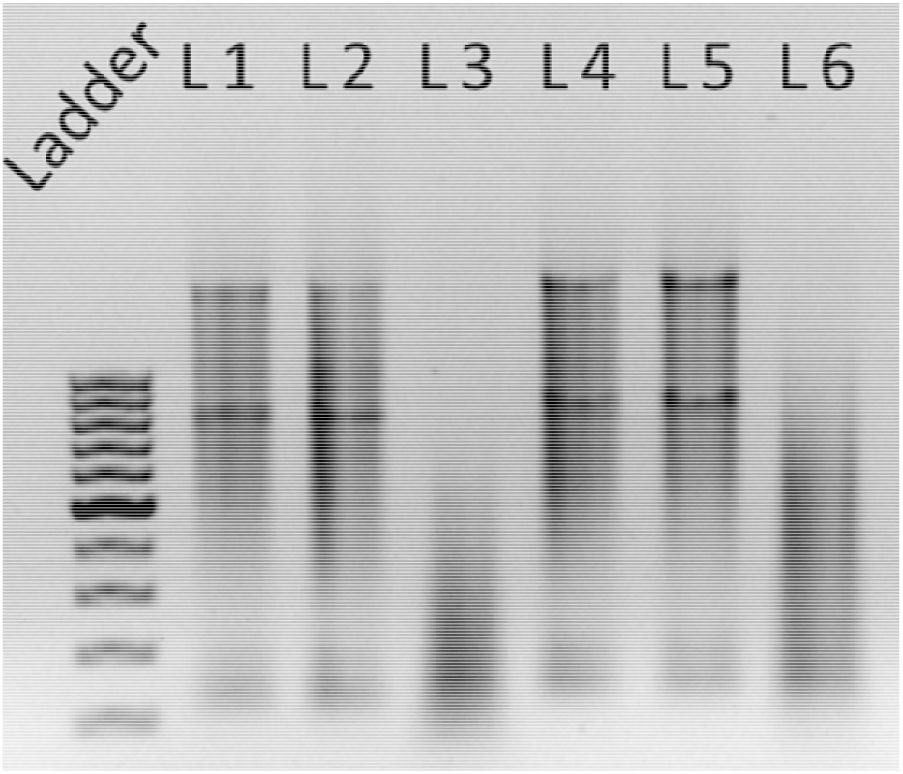
Electrophoretic analysis of RNA samples. It is interpreted from the image that L1, L2, L4, and L5 samples gave positive results for the presence of RNA.

**Figure 7:**
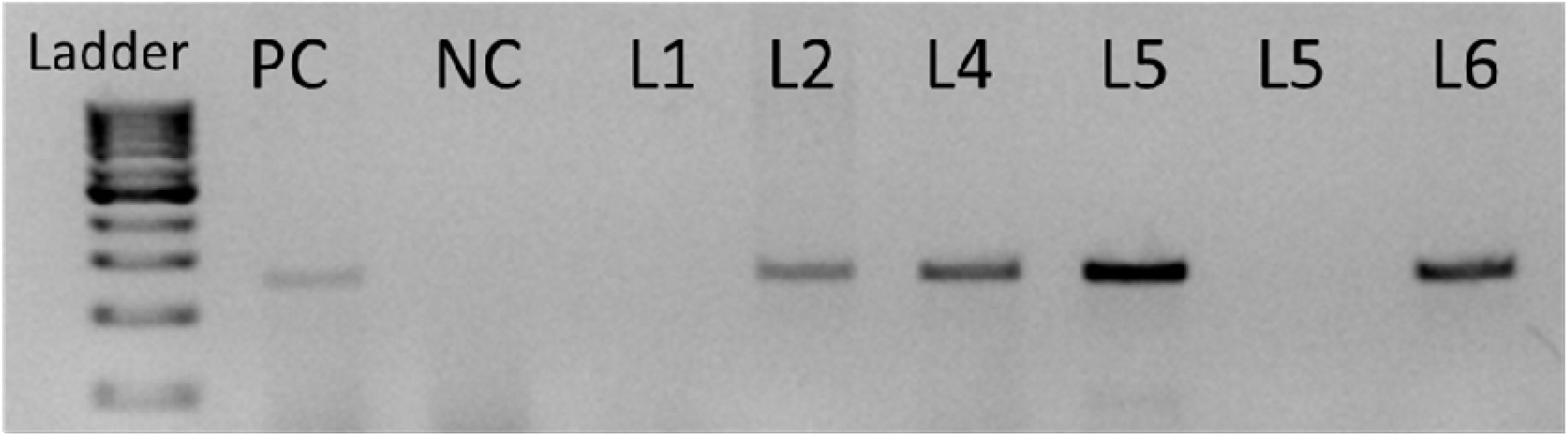
Electrophoretic analysis using cDNA as the template, amplification of the actin housekeeping gene yielded a clear positive band. L1-100bp ladder (Promega), PC-Positive control, NC-Negative control, L2, L4, and L5 showed amplification similar to PC.

## Conclusion

In this study, we developed a novel microfluidic paper-plastic cassette device that integrates a paper substrate for nucleic acid capture and release. This device aids in on-chip nucleic acid extraction. The purity of nucleic acids was analyzed using a Qubit fluorimeter, gel electrophoresis, and PCR amplification. The results were compared with those of the control samples extracted using standard commercially available techniques. This device can extract nucleic acids from biological sources, provided that slight alterations are made to the sample processing before extraction. In summary, the unique features of this technology include rapid DNA extraction, low reagent volumes, extraction of nucleic acids from multiple sources, and easy integration of samples. The ability of paper-based devices to entrap and extract nucleic acids has been extensively studied.

## Ethical Compliance

All procedures performed in studies involving human participants were in accordance with the ethical standards of the institutional and/or national research committee and with the 1964 Helsinki Declaration and its later amendments or comparable ethical standards.

## Data availability

No data was used for the research described in the article.

## Author contribution

**SG**-Experiments design, experimentation, data analysis, manuscript preparation, **UB**-experimentation, data analysis, validation of result and review of manuscript, **SC**-data analysis, validation, review & editing of manuscript, **SK**-result validation, experimental design, **AS**-data analysis, and designing of validation experiments, **VB**-data analysis, validation of result and review of manuscript, **IB**-Methodology design, experimental design, analysis of results, review and editing of manuscript, **AA**-Conceptualization of idea, design of experiments, supervision of experiment, coordination of experimentation and validation process, data analysis, manuscript review, editing and corrections.

